# An adapted typology of tree-related microhabitats including tropical forests

**DOI:** 10.1101/2024.04.20.590405

**Authors:** Ronja Nußer, Giovanni Bianco, Daniel Kraus, Laurent Larrieu, Heike Feldhaar, Matthias Schleuning, Jörg Müller

**Affiliations:** Field Station Fabrikschleichach, Department of Animal Ecology and Tropical Biology, Biocenter, University of Würzburg, Glashüttenstraße 5, 96181 Rauhenebrach, Germany; Senckenberg Biodiversity and Climate Research Centre (SBiK-F), Senckenberganlage 25, 60325 Frankfurt am Main, Germany; Faculty of Biological Sciences, Goethe University Frankfurt, Max-von-Laue-Str. 9, 60438 Frankfurt am Main, Germany; Universitätsforstamt, University of Würzburg, Forstmeisterweg 1, 97437 Haßfurt, Germany; Albert-Ludwigs-Universität Freiburg, Faculty of Environment and Natural Resources, Chair of Silviculture, Tennenbacherstraße 4, Freiburg im Breisgau 79085, Germany; University of Toulouse, INRAE, UMR DYNAFOR, Toulouse, France; Xx CNPF-CRPF Occitanie, Toulouse, France; Animal Ecology I, Bayreuth Center for Ecology and Environmental Research (Bay-CEER), University of Bayreuth, 95440 Bayreuth, Germany; Conservation and Research Department at the Bavarian Forest National Park, Freyunger str. 2, 94481 Grafenau, Germany

**Author notes:** shared last author (Matthias Schleuning, Jörg Müller).

**Keywords:** Tree-related microhabitats (TreMs), Typology, Forest ecosystems, temperate forest, tropical forest

## Abstract

Tree-related microhabitats (TreMs) describe the microhabitats that a tree can provide for a multitude of other taxonomic groups and have been proposed as an important indicator for forest biodiversity (Asbeck et al., 2021). So far, the focus of TreM studies has been on temperate forests, although many trees in the tropics harbour exceptionally high numbers of TreMs. In this study, TreMs in the lowland tropical forests of the Choco (Ecuador) and in the mountain tropical forests of Mount Kilimanjaro (Tanzania) were surveyed. Our results extend the existing typology of TreMs of Larrieu et al. (2018) to include tropical forests and enabled a comparison of the relative recordings and diversity of TreMs between tropical and temperate forests. A new TreM form, Root formations, and three new TreM groups, concavities build by fruits or leaves, dendrotelms, and root formations, were established. In total, 15 new TreM types in five different TreM groups were specified. The relative recordings of most TreMs were similar between tropical and temperate forests. However, *ivy and lianas,* and *ferns* were more common in the lowland rainforest than in temperate forests, and *bark microsoil*, *limb breakage,* and *foliose and fruticose lichens* in tropical montane forest than in lowland rainforest. Mountain tropical forests hosted the highest diversity for common and dominant TreM types, and lowland tropical forest the highest diversity for rare TreMs. Our extended typology of tree-related microhabitats can support studies of forest-dwelling biodiversity in tropical forests. Specifically, given the ongoing threat to tropical forests, TreMs can serve as an additional tool allowing rapid assessments of biodiversity in these hyperdiverse ecosystems.

## Introduction

Forests form terrestrial biodiversity hotspots (Sayer et al., 2019) and provide important supportive (e.g. nutrition, food, and medicine), regulatory (e.g. climate regulation, water, and air purification) and cultural (e.g. education and recreation) ecosystem services (Brockerhoff et al., 2017). However, between 2010 and 2020, annual net global forest loss reached 4.7 million hectares (FAO, 2020), with the greatest total forest loss occurring in the tropics, including 32% of the global loss of forest cover between 2000 and 2012 (Hansen et al., 2013). The main driver of global forest loss is deforestation due to permanent land use changes that favour commodity production and extraction, such as beef, soy, palm oil and wood fibre (Curtis et al., 2018; Laso Bayas et al., 2022). As the loss of forest cover and biodiversity results in a loss of ecosystem services (Reygadas et al., 2023), measures aimed at protecting and restoring forest areas to maintain their provision are needed.

Monitoring whole species communities to evaluate the decline or successful restoration of forest ecosystems is often time-consuming and expensive (Asbeck et al., 2021; Winter et al., 2008), and effective biodiversity indicators are therefore needed (Asbeck et al., 2021; Kozák et al., 2018; Larrieu et al., 2018). In forests, examples of such indicators are stand structure, tree species composition, tree age, diameter distribution, tree regeneration, and deadwood amount and diversity (Larrieu et al., 2018; Larsson et al., 2001). A more recent approach has emerged from the concept of tree-related microhabitats (TreMs), based on the observation that trees provide important microhabitats for a wide range of organisms, including insects, vertebrates, arachnids, Diplopoda, collemboles, gastropods, fungi, lichens, bryophytes, vascular plants, nematodes, rotifers and tardigrades (Bütler et al., 2020; Larrieu et al., 2018; Majdi et al., 2024; Martin et al., 2022; Müller et al., 2014; Schauer et al., 2018; Winter and Möller, 2008). TreMs are morphological singularities found on living or standing dead trees and form essential substrates or microhabitats for thousands of species. Assessments of TreMs can be used to quantify biodiversity at a structural level, by describing habitat quantity (abundance) and quality (diversity).

Kraus et al. (2016) and then Larrieu et al. (2018) compiled a TreM catalogue for temperate and Mediterranean forests, comprising 47 TreM types assigned to 15 groups of 7 main forms according to their morphology and associated taxa (Supplementary Table 1). By using this standardized catalogue, several studies have reported a negative effect of forest management intensity on the abundance of most TreMs and on TreM diversity (e.g., Paillet et al., 2017; Winter and Möller, 2008). For example, broadleaf trees as well as larger, senescent, or dead trees carry more TreM types such that TreM diversity is higher in natural than in managed forests (Courbaud et al., 2022; Larrieu et al., 2018; Martin et al., 2022). Moreover, TreM composition is sensitive to the type of forest management, with specific TreMs, such as *dendrotelms* and *bark-loss*, being promoted by logging (Larrieu et al., 2012; Vuidot et al., 2011).

So far, most studies of TreM abundance and diversity have been conducted in temperate forests (Martin et al., 2022). For tropical forests, biodiversity indicators still need to be developed in order to assess the changes in their diversity. In contrast to temperate regions, the tropics have retained many of their old-growth forests (Sabatini et al., 2021), which potentially host a huge diversity of TreMs. Due to the rapid growth and high species diversity of trees in tropical forests (Brandon, 2014), their TreMs are likely to be more diverse than those in temperate forests. In addition, large areas of the tropics contain secondary forests, but little is known about their impact on biodiversity (Brandon, 2014; Chazdon, 2014). The scarcity of knowledge of TreM composition and diversity in tropical primary and secondary forests is at least in part due to the lack of an adapted TreM typology for tropical forests.

Thus, in this study we systematically sampled TreMs in two tropical regions, one in Africa and the other in South America, differing in their tropical forest ecosystem types. Our objectives were (1) to expand and adapt the catalogue of TreMs to tropical forests and (2) to compare the abundance and diversity of TreMs at the tree level among different forest ecosystems.

## Materials and Methods

### Study sites

The first tropical TreM assessment was conducted on the plots of the REASSEMBLY project (FOR 5207) (www.reassembly.de) of the Choco, in Canandé, Ecuador (0.56° N, –79.20° W), part of the large Choco ecosystem. The project consists of 62 plots (50x50 m) along a recovery gradient ranging from active pastures and cacao plantations, recovering forests ranging in age from 2 to 38 years and old-growth forests. No active planting has taken place on the recovery plots. All plots are located between 100 m and 600 m above sea level (Escobar et al., 2024).

The second tropical study site comprised 65 study plots on the southern slopes of Mount Kilimanjaro, Tanzania (3.00° S, 37.22° E). The plots were established as part of the DFG research units KiLi (FOR 1246) and Kili-SES (FOR 5064). Mount Kilimanjaro is a freestanding dormant volcano, rising from a plateau at 700 m above mean sea level (AMSL) to its summit at 5895 m AMSL. Along this altitudinal gradient lies a broad diversity of ecosystems, from dry savannahs in the lowlands to montane forests in the rain belt and afro-alpine meadows at the highest elevations. At Mount Kilimanjaro, TreM abundance and diversity were assessed in the 46 tree-containing plots (50 × 50 m in size). These plots are distributed alongside five elevational transects and represent ten key ecosystem types (wild savannah, culti-vated maize, home gardens, coffee plantations, lower montane forest, *Ocotea* forest, disturbed *Ocotea* forest, *Podocarpus* forest, disturbed *Podocarpus* forest, *Erica* forest) (Hemp, 2006).

The relative recordings of TreM types in the two tropical regions were compared using recently published, comprehensive data from temperate beech forests, as in temperate forests this forest type typically harbours the largest number of TreMs (Courbaud et al., 2022). The beech forests considered in this study included both managed and old-growth forests located in France, Switzerland, Germany, Bulgaria, Ukraine, Georgia, Armenia, and Iran (Mamadashvili et al., 2023), thus covering the entire temperate beech belt in the western Palaearctic.

### TreM surveys

In Ecuador, up to 30 trees in each of the 62 plots were surveyed from August to October 2023. In some plots, particularly those in pastoral areas, the sparse presence of remnant trees resulted in the observation of fewer trees; thus, 25 trees per plot were surveyed on average. Trees were selected using a relascope [Spiegelrelaskop® (22823), after W. Bitterlich], which uses angle-count sampling. All trees with a basal area factor (≥ 3were marked around three randomly selected points per plot. If this did not result in 10 trees per sampling point, the count size was reduced until enough trees had been marked. The angle-count sampling method (Bitterlich 1984) was chosen as it was previously used to obtain data from temperate forests (Mamadashvili et al., 2023). The focus of this method is to sample a balanced number of trees in all dimension classes. It grants a greater likelihood of selection to larger trees. The selected trees were inspected from all sides and binoculars were used to identify TreMs in the tree crown.

On Mt Kilimanjaro, TreM surveys took place on 48 plots. The five largest trees were selected in each plot. If trees were similarly sized, the sampling of trees of different species was priori-tised. For trees taller than 5 m, a rope-based technique was employed to access the tree canopy. Once the highest anchor point had been reached, the tree climber proceeded to descend the entirety of the tree, pausing at 1-m intervals to assess the presence of each microhabitat type within that interval.

### TreM classification

The main objective of this study was to extend the typology of Larrieu et al. (2018) to include tropical forests. According to Larrieu et al. (2018), a TreM is a habitat resource formed by a singularity (e.g. rockfall, skidder damage), regardless of its trigger. Different TreMs can be identified when there are differences between the assemblages of their associated taxa. During TreM surveys in the lowland tropical rainforest of Choco, new TreM types were identified and assigned to the existing typology. On Mt. Kilimanjaro, the TreM survey also followed the catalogue developed by Larrieu et al. (2018), with the following exceptions: (1) the wood-pecker cavities recorded at Mt. Kilimanjaro were assigned to the TreM type *large woodpecker breeding cavity* (TCV13; (2) the fruiting bodies of saproxylic fungi were not distinguished from those of slime moulds; rather, a distinction was made only between polypores and all other fungi, corresponding to the TreM types *perennial polypores* (TFB11) and *pulpy aegaric* (TFB13). All newly found and described TreM types are presented in the Results (Table *1*).

**Table 1:**
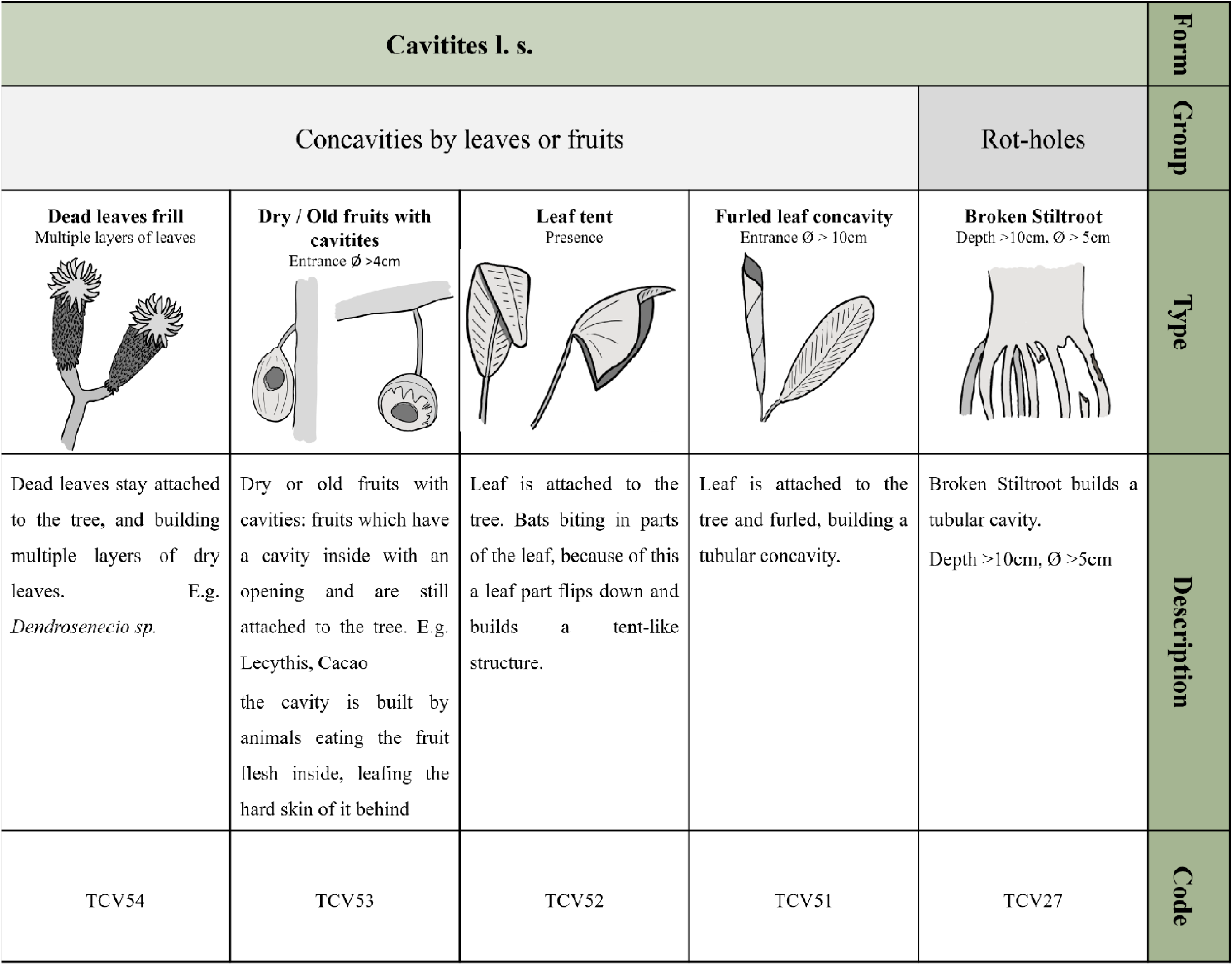
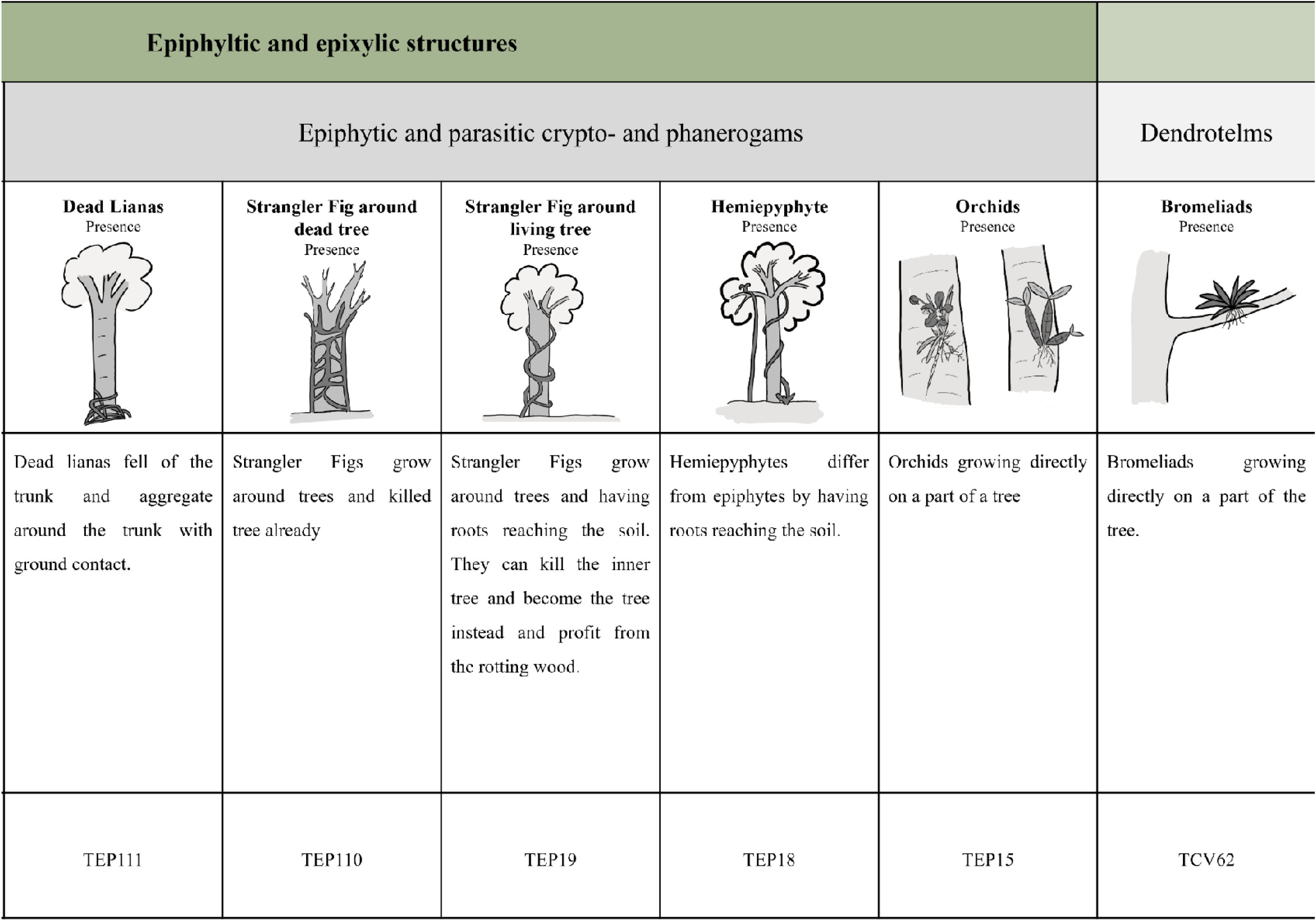
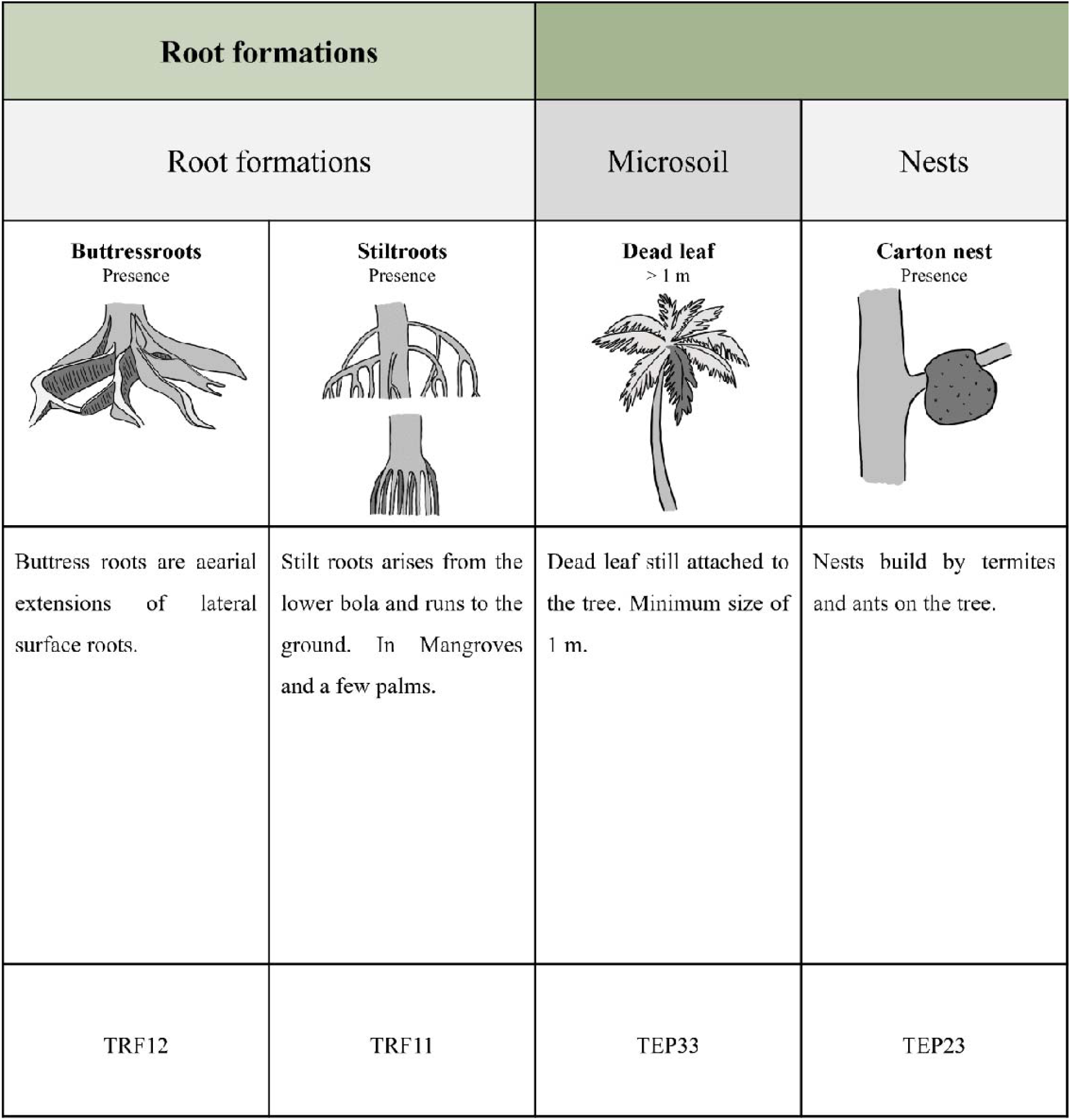
New TreM types from the catalogue of tree-related microhabitats in tropical forests. TreMs are visualised with drawings, a short description and a proposed size threshold. To facilitate field surveys, each TreM was assigned a code, composed of the letter T for tropics followed by two letters standing for the TreM form (e.g. “CV” for “Cavities”). The next number stands for the TreM group and the last number for the TreM type.

### Data analysis

The diversity and abundance (relative recordings) of TreMs at the tree level between different forest ecosystems were compared based on the frequency of each TreM type for each forest ecosystem. The relative recordings of TreMs and their diversity were then calculated using rarefaction-extrapolation curves, plotting the recordings along Hill numbers (1973) (effective numbers of species). Hill numbers have been proposed as a generalised approach to biodiversity measurement (Ellison, 2010), as they unify numerous indices along a continuous gradient of weighted abundances. In this study, Hill numbers were used to control for differences in sampling effort among three different data sets, representing the total number of species (q = 0), Shannon’s entropy (q = 1) and Simpsons’s index (q = 2) (Hill, 1973). Hill numbers can also be related to species composition, as the three numbers represent rare, common and dominant species, or in this case TreM types (Chao et al., 2014). All analyses were conducted in R Studio using R version 4.3.2. The packages ggplot2 (Wickham, 2016) and ggthemes were used for visualisation, and the package iNext (Hsieh et al., 2016) for computing diversity measures.

## Results

### New TreMs from tropical forests

One new TreM form, Root formations, and three new TreM groups, concavities build by fruits or leaves, dendrotelms, and root formations. were defined in this study. In addition, 15 new TreM types occurring in tropical forests and not included in the original typology (Table 1) were identified. Our extended typology of Larrieu et al. (2018), including all new tropical TreM types, is presented in Supplementary Table 1.

For the form Cavities *sensu lato*, a new TreM group consisting of concavities built by leaves or fruits was defined. This new group encompassed four different types of TreM: (1) *furled leaves*. consisting of large, incompletely developed leaves that form a tubular cavity with an opening at the top; (2) *leaf tent*, made by bats biting into the midrib of the leaf to create a shelter underneath (Kunz et al., 1994); (3) *dry fruit with cavity*, which refers to the cavity made by animals in the flesh of the fruit, but not the hard skin, resulting in a concavity inside the fruit (entrance Ø >4 cm); (4) *dead leaves frill*, arising from dead leaves that remain on the tree, instead of falling to the ground (e.g. *Dendrosenecio* sp.), such that their layers form small concavities (Table *1*).

To the TreM group rot holes, the type *broken stiltroot* was added. Due to physical damage, stilt roots can break and form a tubular cavity, usually open at the bottom. However, if the root is penetrated (e.g. by fallen trees or branches), it may also be open at the top, with ground contact. If the root is only severely damaged, an opening may form that grants access to a tubular cavity. The proposed threshold to record this TreM is an entrance at least 5 cm in diameter and a concavity at least 10 cm deep. In addition, we defined the TreM group dendrotelms, which contains the TreM type *dendrotelm* from the original catalogue along with our newly added type *bromeliads* (Bromeliaceae) (Table *1*).

Seven TreM types were added to the form of epiphytic and epixylic structures, five of which belong to the group epiphytic and parasitic crypto- and phanerogams., e.g. epiphytic *orchids* (Orchidaceae) and *hemiepiphytes* growing on trees. These types differ from epiphytes in that their roots reach down to the ground, with some growing around the tree, such that a climbing trunk and a crown are formed (e.g. strangler figs), sometimes killing the host tree and resulting in the accumulation of its decaying wood (Athreya, 1997). These strangling hemiepiphytes were added as two separate TreM types, with strangling hemiepiphytes around a living tree with net-like growing climbing organs (i.e. *strangler fig around living tree*) distinguished from strangling hemiepiphytes that already killed the inner tree (i.e. *strangler fig around dead tree*), since the latter also contain saproxylic substrates. In addition, a new TreM type, *dead lianas*, was defined, which refers to dead lianas around the trunk that are connected to the soil or root system of the tree (Table *1*). The TreM type *carton nests*, built by termites (Termitoidae) or ants (Formicidae) out of wood fibre and faecal material on the trunk or a branch of a tree, was separated from the type *invertebrate nests* (e.g. caterpillar nests). They can be quite large in the tropics (Collins et al., 1997; Haverty et al., 1997) and have a higher stability (Andrews, 1911; Emerson, 1938; Hubbard, 1877; Lubin and Montgomery, 1981; Thorne, 1980) than in other climate zones. Within the TreM group microsoil, the type *dead leaf* was defined as large dead and decaying leaves (> 1 m in size) still attached to the tree. The different types of roots found in the tropics were covered by adding the TreM type root formation, divided into two types, *stiltroots* and *buttressroots.* Not all *buttressroots* contain concavities, which are already described by the TreM type *buttress root concavity;* rather this type refers to a microhabitat that hosts other tropical taxonomic groups (Table *1*).

### Diversity and compositional changes in TreMs

In the lowland tropical rainforest of Canandé (Ecuador), 1573 trees were sampled and 57 TreM types were found, including 14 of the new TreM types. In total, 5060 TreMs were recorded. In mountain tropical forests (Tanzania), the 180 sampled trees harboured 42 TreM types, with 1537 recordings. In temperate forests, the 4506 sampled trees harboured 45 different TreM, with 6116 recordings.

The most common TreM types in the lowland tropical rainforest were *ivy and lianas* (TEP13) accounting for 18% of all recordings, followed by *hemiepiphytes* (TEP18) and *bromeliads* (TCV62), accounting for 10% and 9% respectively (Figure 1). In the mountain tropical forests, the most common TreMs were *dead branches* (10%), followed by *foliose and fruticose lichens* (TEP12) and *bryophytes* (8% each) (Figure 1). The TreMs *foliose and fruticose lichens*, *bark microsoil* (TEP31) and *limb breakage* (TIN22) were more common in mountain tropical forests than in lowland forests. In the temperate forest, the most common TreM types were *dead branches* (TDE11), *bryophytes* (TEP11) and *trunk base rot-holes* (TCV21), comprising 19%, 15% and 8% of all records respectively (Figure 1). *Ivy and lianas and ferns* (TEP14) were more common in tropical rainforest, and *bryophytes* and *dead branches* in the temperate forests. The differences between the mountain tropical forests of Mt. Kilimanjaro and the lowland rainforest of Canandé included the more common occurrence of *foliose and fruticose lichens*, *bark microsoil* (TEP31) and *limb breakage* (TIN22) in the former.

**Figure 1:**
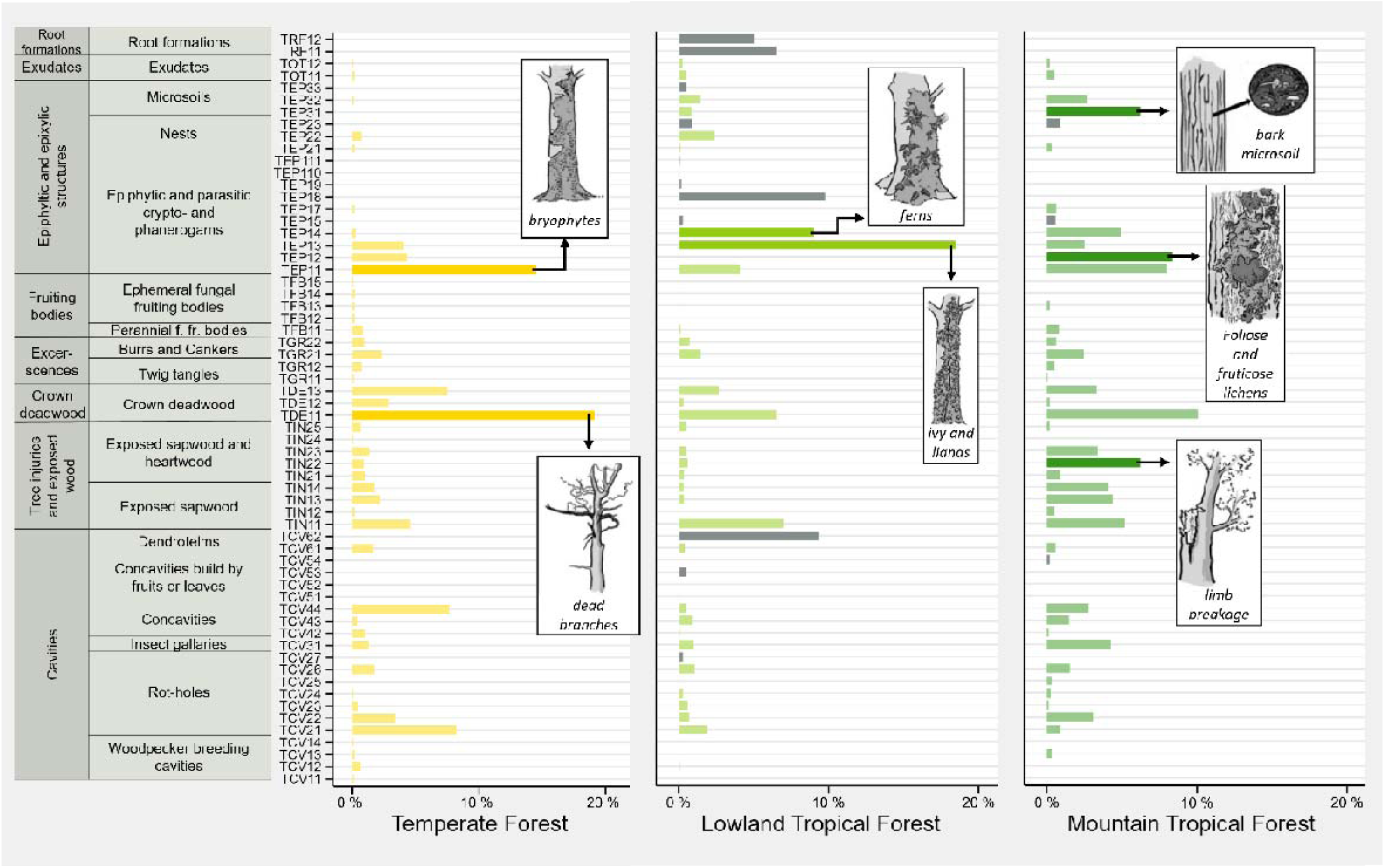
Comparison of tree-related microhabitats (TreM) in temperate beech-dominated forests with a tropical lowland rainforest in Canandé, Ecuador, and the tropical montane forests of Mt. Kilimanjaro, Tanzania. The presence of each TreM type is shown as a percentage of all recordings for each region. Larger differences are indicated by a darker colour and by drawings of the TreM types. New TreM types are shown in grey.

### Comparison among forest ecosystems

Diversity analyses showed that TreM diversity, with a focus on rare TreM types, was highest in the lowland tropical forest, suggesting that TreM richness was highest in this forest ecosystem. By contrast, the mountain tropical forest hosted the largest TreM diversity with respect to common (q = 1) and dominant (q = 2) TreM types. While temperate forests hosted the lowest diversity of TreMs for common and dominant species, their richness in rare TreM types was similar to that of the mountain tropical forest. The curves indicated that the sampling of common and dominant species was almost complete for all forest ecosystems, whereas rare species of lowland tropical forest were estimated to have been incompletely sampled (Figure 2).

**Figure 2:**
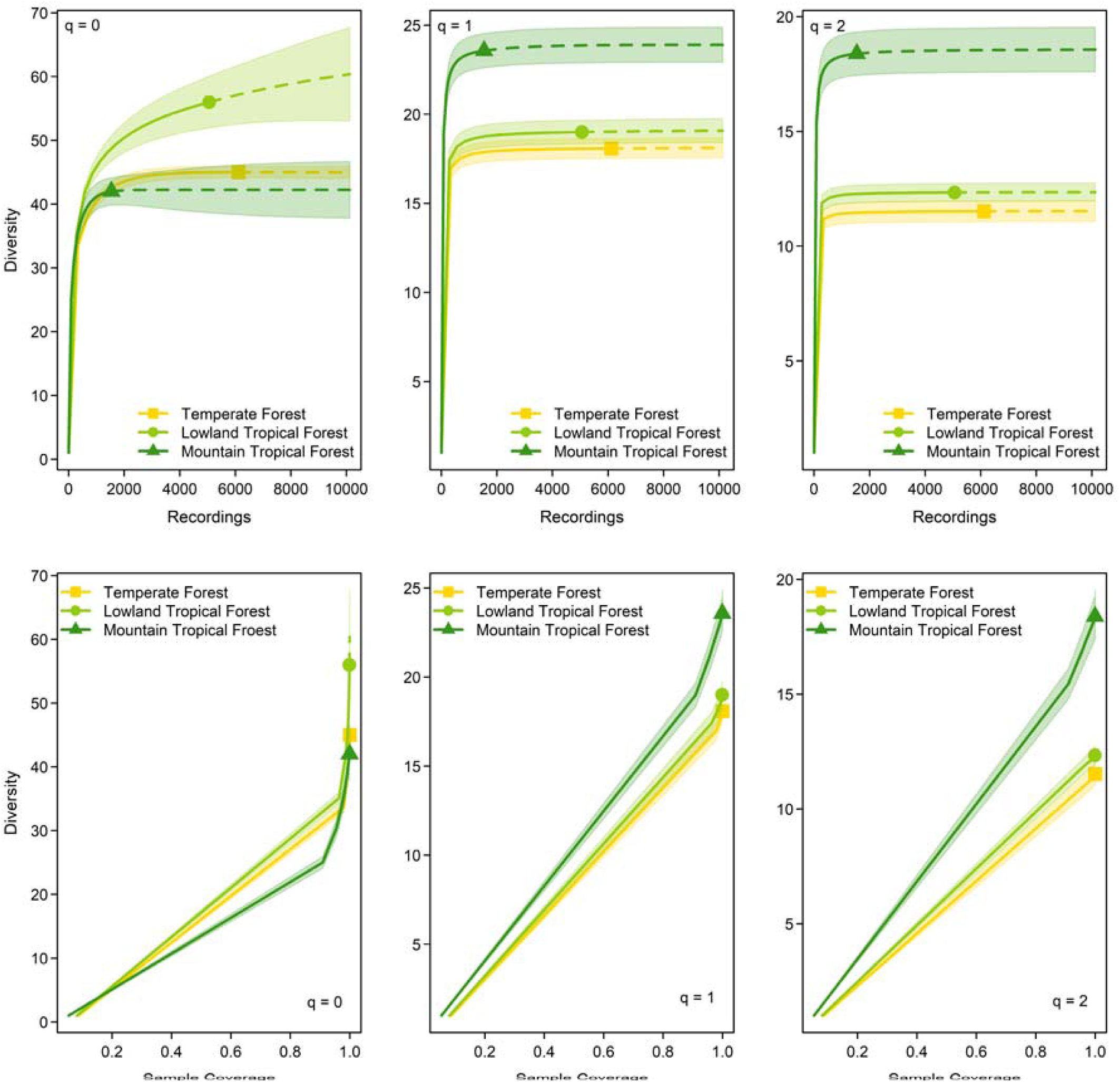
Diversity estimates along the Hill numbers with respect to sample size and sample coverage for the temperate forest, lowland tropical forest and mountain tropical forest. The total number of recordings of different TreM-types is shown for each forest type in relation to TreM diversity. The Hill numbers can be interpreted as rare (q = 0), common (q = 1) and dominant (q = 2) TreM types. Sample coverage of each forest ecosystem is shown as the coverage based rarefaction/extrapolation curve along the Hill numbers.

## Discussion

Our study resulted in the definition of one new TreM form (Root formations) and three new TreM groups (concavities build by fruits or leaves, dendrotelms, and root formations). In addition, 15 new TreM types were identified in tropical forests that were not included in the original catalogue, which was based only on observations in temperate and Mediterranean forests (Table *1*). The observed differences in TreM composition and diversity between the three forest ecosystems suggested that tropical forests host a larger overall diversity of TreMs than temperate forests. While we are aware that our survey was limited to two tropical forests and did not follow a fully standardised protocol at the two sites, the resulting TreM typology for tropical forests provides a benchmark for future studies in tropical forests.

### New TreM types from tropical forests

Due to their different structures, substrates and microclimatic conditions, the newly described TreM types host different animal species. Biotic and abiotic factors play an important role in cavity formation, which makes each cavity unique (Henneberg et al., 2021; Micó, 2018; Schauer et al., 2018). Thus, it is possible that, for example, *broken stiltroots*, with their tubular cavity close to the ground, host an insect fauna distinct from that of the rot holes of temperate forests.

The newly described TreM type *furled leaf concavity* provides a specific microclimate, preferred, for example, by leaf-roosting bats (Pérez-Cárdenas et al., 2019). Bats, such as the Honduran white bat (*Ectophylla alba*), build tents from leaves by biting into the midrib (Rodríguez-Herrera et al., 2006), creating a unique TreM. These *leaf tents* support their mating system but are also used as roosts (Kunz et al., 1994). The temperatures under leaf tents can be higher than in the surrounding area, especially at night, which could reduce energy costs for bats (Rodríguez-Herrera et al., 2016). The TreM type *dry fruits with cavities* could be used as shelters by arthropods. The shells of large fruits may form container-like habitats that hold rainwater. However, if the soft parts remain and rot, the liquid will be extremely eutrophic (Yule and Sen, 2004) and likely be used by mosquitoes as a breeding substrate (Machado-Allison et al., 1988).

In the tropical montane forests, the new TreM type *dead leaves frill* was found on trees of the genus *Dendrosenecio*. These trees retain a dense frill of dead leaves that insulates the pith cells and helps the plant to cope with the stark changes in temperature that characterise afroalpine meadows (Beck, 1986). The presence of various species of arthropods on these *dead leaves frills* suggested that small animals burrow within the dead leaves to withstand temperature changes.

In bromeliads, temporary pools of water (phytotelms) form a microhabitat for a large number of organisms, some of which have become specialised for it (Kitching, 2001). While the outer, more mature leaves hold separate water bodies, the younger, inner leaf axils combine to form a common basin (Kitching, 2001). These structures allow the colonisation of many amphibian species (Peixoto, 1995; Silva et al., 2011), providing shelter for some but needed throughout the life cycle, including for reproduction and feeding, for others, such as dendrobatid species (Anura, Dendrobatidae), which breed in bromeliads (Masche et al., 2010; Savage, 2002; Twomey and Brown, 2009).

Some *orchids* (e.g. *Caularthron bilamellatum*) produce pseudobulbs that develop a central cavity with age. A number of ant species use these cavities to build nests (Dutra and Wetterer, 2008; Fisher, 1992; Yanoviak et al., 2011). We also described several *hemiepiphytes* growing on and around trees. Their growth structure, especially that of the strangler fig, can create small shady, moist habitats. Some *hemiepiphytes* secrete a thick layer of mucilage on their aerial roots to limit desiccation and protect against pathogens, thus providing a favourable habitat for larvae of the psychodid fly *Mucomvia emersa* (Kvifte et al., 2018), for example. *Hemiepyphytes* are also a source of food for birds and some mammals (Arren and Brockelman, 1982).

Another TreM structure specific to the tropics consists of the *carton nests* build by termites (Termitoidae) or ants (Formicidae). Because of their special microclimatic conditions, unique structure and ecological importance in the tropical forests, these were separated from other *invertebrate nests*, such as galls, and comprise a new TreM type in the TreM group nests. However, extending the TreM types *invertebrate nests* and *carton nests* in size and form might be required in the future to capture the larger variety presented in the tropics. The microclimate in *carton nests* is warmer, more humid and less variable than ambient conditions (Dechmann et al., 2004; Fuller and Postava-Davignon, 2014). Moreover, this microhabitat is used by a variety of organisms, including some vertebrates. For example bats (e.g. *Lophostoma silvicolum*) (Dechmann et al., 2004) as well as some species of birds inhabit the active arboreal nests of *Nasutitermes cortiger* (Kesler and Haig, 2005; Sazima and D’Angelo, 2015). Arboreal termite nests are also often used by ants for foraging and/or nesting (Santos et al., 2010). Among the organisms using ant-made *Carton nests* is the gecko *Gonatodes humeralis,* which selects arboreal *Azteca* nests as oviposition sites (Gotwald, 1984). Beyond animals, *carton nests* provide a suitable substrate for the growth of epiphytes, which may also be colonised by the host ant colony (Davidson and Epstein, 1989). In building their carton nest, ants incorporate the seeds or fruits of certain epiphytic plants while the plant roots confer stability to the nest carton (Davidson and Epstein, 1989; Kaufmann and Maschwitz, 2006).

Finally, *buttress roots* are specific to tropical forests and their positive impact on the abundance and species richness of herpetofauna, due to the accumulation of leaf litter and the high humidity, has been described (Sutton and Memmott, 1994; Whitfield and Pierce, 2005). They also support a higher abundance and richness of adult and larval macroinvertebrates in the leaf litter (Alencar et al., 2011; Somerville, 2010).

### Compositional changes in TreMs

In the lowland tropical rainforest, the TreMs *ivy and lianas* and *hemiepiphytes* and *bromeliads*, and in temperate forests the TreMs *dead branches*, *bryophytes,* and *trunk base rot-holes* were the most common. The main differences between these two regions forest types were the proportions of *ivy and lianas* and *ferns*, both of which were more dominant in the lowland tropical forest, and the proportions of *bryophytes* and *dead branches*, which were abundantin temperate forests. Most epiphytic structures were rare in temperate forests (except *bryophytes*). A study comparing tropical and temperate plant growth forms and their vertical distribution showed that about 20% of epiphytic species are found in tropical forests, and only 1% in temperate forests (Spicer et al., 2020). Similarly, *lianas* are more widespread in the tropics, and in tropical lowlands are considered indicators of disturbed forests under warmer temperatures with low amounts of precipitation (Ngute et al., 2024). Bryophytes, by contrast, are more common in temperate forests. While they are highly dependent on external water, they are also sensitive to high temperatures and to drought (He et al., 2016; Zotz and Bader, 2009).

Trunk rot holes were more abundant in temperate forests than in the tropical lowland forest. The formation of *rot holes* is promoted by factors such as tree age, tree species and environmental conditions, including direct sun exposure, poor soil and low precipitation (Larrieu et al., 2022). With increasing age, the probability that the tree will be wounded and infected, resulting in decay and the formation of rot holes, increases. Both the development stage and the tree species influence the compartmentalisation capacity of a tree. A low capacity enhances fungal colonisation and decay, leading to rot holes. The compartmentalisation capacity decreases as tree development proceeds (Smith, 2015). The high species richness would lead to more trees with a high compartmentalization capacity. However, some tree species in tropical forests are very efficient in wound closure, to avoid infections (Morris et al., 2016; Turner, 2001), which would reduce the formation of rot holes.

In the montane tropical forests, *dead branches*, *foliose and fruticose lichens* and *bryophytes* were the most common TreM types. *Lichens* are often favoured by moist climate at higher altitudes (Asbeck et al., 2019; Larrieu et al., 2022). Differences in the proportions of *bark microsoil,* and *limb breakage* between the lowland and the montane tropical forests were also determined. The higher abundance of *limb breakage* in the mountain tropics can be explained by the increase in wind speed with altitude (Oliver, 1971), with strong winds causing *limb breakage* (Larrieu et al., 2022). *Bark microsoil* is formed by mosses, lichens or epiphytic alga residues and decaying bark (Bütler et al., 2020). The low abundance of *bryophytes* in the lowland rainforest and the high abundance of *lichens* in the mountain region could explain why *bark microsoils* were more common in mountain tropical forests.

### Diversity differences in TreMs

The highest diversity of common and dominant TreM types was detected in the mountain tropical forest, whereas rare TreM types, and thus TreM richness, were more frequent in the lowland tropical forest. Thus, while TreMs were a frequent feature of the mountain tropical forest, their overall richness tended to be lower than in the tropical lowland forests. The reasons for these differences could be specific climate conditions, such as those promoting lichen-associated microhabitats at high altitudes (Bässler et al., 2016), but also the lower tree species diversity in mountain than in lowland tropical forests. Although the differences associated with the variation in tree species richness between lowland and montane forests may have been further accentuated by the larger number of tree individuals sampled in the lowlands, it is still plausible that the highly diverse tropical lowland forests host the largest TreM richness globally.

Overall, the lowest diversity of common and dominant TreM types was found in temperate forests, where altitude, management and DBH are the main drivers of TreM diversity (Asbeck et al., 2019; Larrieu and Cabanettes, 2012; Mamadashvili et al., 2023; Winter and Möller, 2008). However, as these drivers vary within the studied areas, they are unlikely to explain the differences in diversity among ecosystems. It is more likely that the higher biodiversity in tropical than temperate forests is reflected in the diversity differences of TreMs. Such latitudinal differences in biodiversity from equatorial to polar regions have been reported not only for organisms but also for habitat diversity (Hillebrand, 2004; Mannion et al., 2014). Important causes for this almost universal pattern have been attributed to energy availability, which is higher in the tropics (Turner, 2004), and to higher environmental heterogeneity in tropical than temperate ecosystems (Ricklefs, 1977). A more comprehensive understanding of the global patterns of TreM diversity awaits further comparative studies in more tropical forests using the expanded TreM catalogue.

## Conclusion

Our survey of TreMs in tropical forests revealed new TreMs, including those specific to tropical forests, and important differences in TreM diversity and composition among forest ecosystems. The expansion of the TreM typology to include TreMs in tropical forests will facilitate further surveys of these microhabitats in tropical forests and open up new avenues for future research. For instance, the proposed tropical TreM catalogue could be applied to assess the ecological integrity of old-growth tropical forests or the success and pace of recovery in tropical forests. As TreMs play essential roles in the survival and reproduction of a diverse set of taxa, they could also serve as an important indirect indicator of tropical biodiversity.

## Acknowledgments

This work was funded by the Deutsche Forschungsgemeinschaft (DFG) within the Research Units REASSEMBLY (FOR 5207) and Kili-SES (FOR 5064). We would like to thank the REASSEMBLY group for their logistic support and are indebted to our research assistants Miguel Angel Tacuri Chichande and Alexis Leonardo De La Cruz Aveiga for their help in the field. We also thank the Fundación Jocotoco and Fundación Tesoro Escondido for logistic support and permission to do research on their reserves. We thank the COSTECH, TAWIRI and TANAPA authorities for providing the permits to conduct fieldwork on Mount Kilimanjaro (permit no: 2022-307-NA-2021-094). We thank the staff at the research station of Nkweseko (Moshi, Tanzania) for hosting us and supporting our fieldwork. Special thanks go to Raymond Vitus, Frederick Issaack and Esrom Nkya who assisted with the TreM survey. Furthermore, we thank Wendy Ran for her professional language revision.

