## Supplementary material for "An adapted typology of tree-related microhabitats including tropical forests": Supplemtal Table 1

Supplement Table 1: Complete typology of tree-related microhabitats in tropical forests. TreMs are visualised with drawings, a short description and a proposed size threshold. To facilitate field surveys, each TreM was assigned a code, composed of the letter T for tropics followed by two letters standing for the TreM form (e.g. “CV” for “Cavities”). The next number stands for the TreM group and the last number for the TreM type.

| **Form** | **Group** | **Type** | **Description** | **Code** |
| --- | --- | --- | --- | --- |
| **Cavitites l. s.** | Woodpecker breeding cavities | **Small woodpecker breeding cavity**  Entrance Ø <4cm  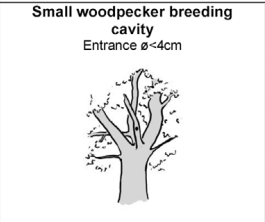 | Cavity entrance ø < 4 cm.  The breeding cavity of small-sized woodpeckers drilled in a dead branch. | TCV11 |
|  |  | 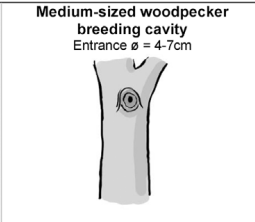**Medium-sized woodpecker breeding cavity**  Entrance Ø = 4-7cm | Round cavity entrance about ø = 4–7 cm.  The breeding cavities of the medium-sized woodpeckers are usually drilled into decaying wood (dead branch, snag, insertion of broken-off branches). | TCV12 |
|  |  | **Large woodpecker breeding cavity**  Entrance Ø >10cm  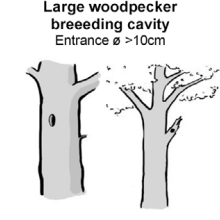 | Oval cavity entrance ø < 10 cm.  The breeding cavities of large-sized woodpeckers are usually drilled on the main part of the trunk (without branches). | TCV13 |
|  |  | **Woodpecker flute**  Entrance Ø > 3cm  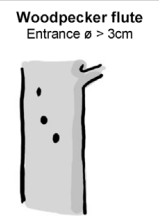 | At least three woodpecker breeding cavities in line on the trunk.  Maximum distance of 2 m between two consecutive cavities. | TCV14 |
|  | Rot-holes | **Trunk base rot-hole (closed top, ground contact)**  Opening Ø > 10cm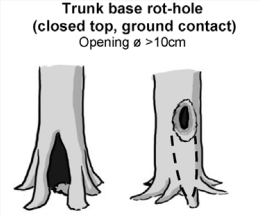 | Cavity chamber is completely protected from surrounding microclimate and rain Top-closed trunk cavity containing more or less mould (depending on its development stage). The cavity bottom has ground contact. Note that the cavity entrance can be higher on the trunk. | TCV21 |
|  |  | **Trunk base rot-hole (closed top, no ground contact)**  Opening Ø > 10cm  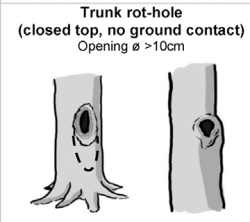 | Top-closed trunk cavity containing more or less mould (depending on its development stage). The cavity bottom has no ground contact. | TCV22 |
|  |  | **Semi-open trunk rot-hole**  Opening Ø > 30cm  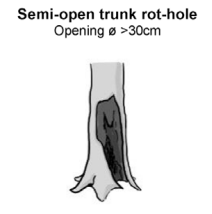 | Cavity chamber is not completely protected from surrounding microclimate and rain may flow in. Note that the cavity entrance can be higher up in the trunk | TCV23 |
|  |  | **Chimney trunk base rot-hole**  Opening Ø > 30cm  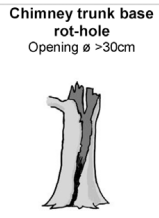 | Cavity in the trunk of the tree that is completely open at the top, often resulting from stem breakage; the cavity base reaches ground level, so the inner cavity is in direct contact with the soil. | TCV24 |
|  |  | **Chimney trunk rot-hole**  Opening Ø > 30cm  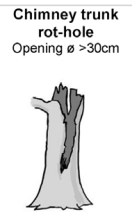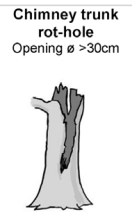 | Cavity in the trunk of the tree that is completely open at the top, often resulting from stem breakage; the cavity base does not reach ground level, so the inner cavity is not in direct contact with the soil. | TCV25 |
|  |  | **Hollow branch**  Opening Ø > 10cm  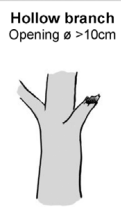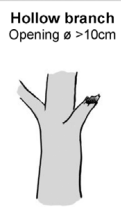 | Rot hole in a large branch, resulting in a tubular shelter, often horizontally oriented | TCV26 |
|  |  | **Broken Stiltroot**  Depth >10cm, Ø > 5cm  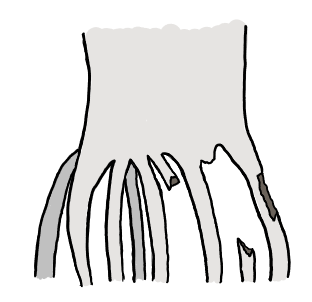  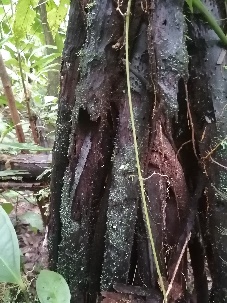 | Broken Stiltroot builds a tubular cavity.  Depth >10cm, Ø >5cm | TCV27 |
|  | Insect galleries | **Insect galleries and bore holes**  Hole Ø > 2cm or area >300cm²  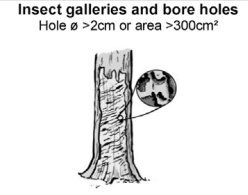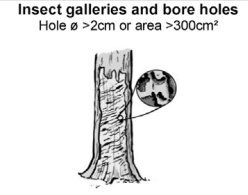 | A bore hole network of xylophagous insects indicates a wood hole system. An insect gallery is a complex system of holes and chambers created by one or more insect species in the wood. | TCV31 |
|  | Concavities | **Woodpecker foraging excavation**  Depth >10cm, Ø > 10cm  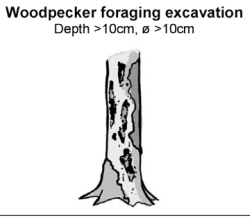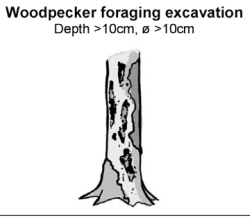 | Concavity resulting from the foraging activities of woodpeckers. The excavation is conical: the entrance is larger than the interior. | TCV42 |
|  |  | **Trunk bark-lined concavity**  Depth >10cm, Ø > 10cm  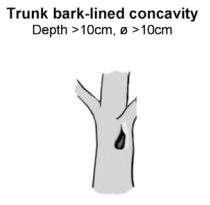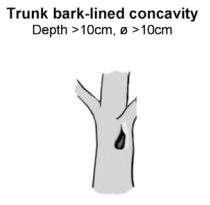 | Natural bark-lined concavity on the tree trunk. No mould. | TCV43 |
|  |  | **Root-buttress concavity**  Entrance Ø > 10cm  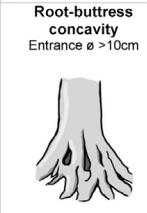 | Natural bark-lined concavity at the base of the tree trunk formed by the tree roots and the soil. No mould (if so: see Trunk base rot hole) | TCV44 |
|  | Concavities by leaves or fruits | **Furled leaf concavity**  Entrance Ø > 10cm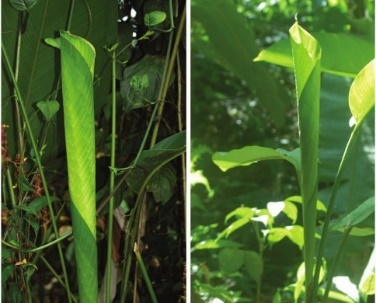  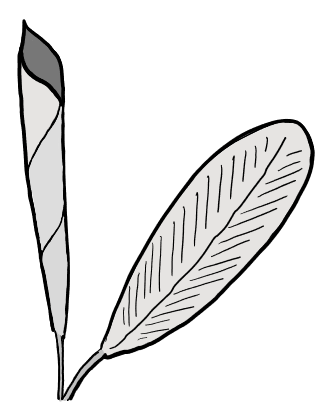 | Leaf is attached to the tree and furled, building a tubular concavity. | TCV51 |
|  |  | **Leaf tent**  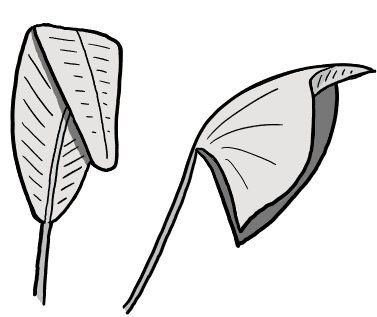 | Leaf is attached to the tree. Bats biting in parts of the leaf, because of this a leaf part flips down and builds a tent-like structure. | TCV52 |
|  |  | **Dry / Old fruits with cavitites**  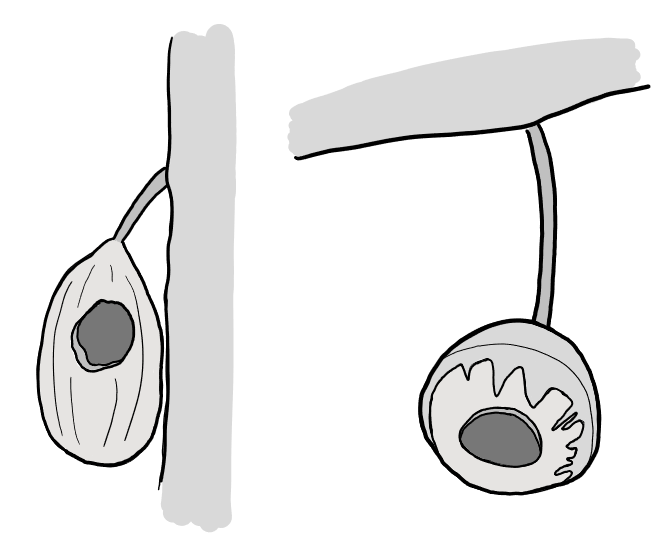 | Dry or old fruits with cavities: fruits which have a cavity inside with an opening and are still attached to the tree. E.g. Lecythis, Cacao  the cavity is built by animals eating the fruit flesh inside, leafing the hard skin of it behind | TCV53 |
|  |  | **Dead leaves frill**  Multiple layers of leaves  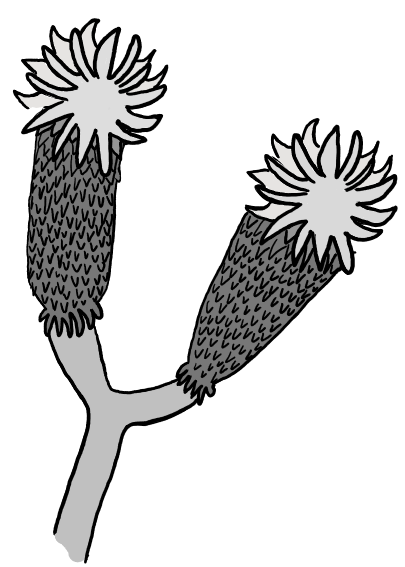 | Dead leaves stay attached to the tree, and building multiple layers of dry leaves. E.g. *Dendrosenecio sp.* | TCV54 |
|  | Dendrotelms | **Dendrotelm**  Ø > 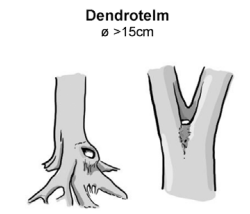15cm  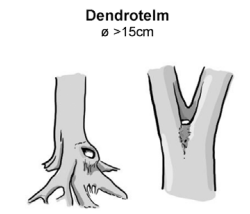 | Cup-shaped concavity that, due to its form, retains water until it dries out by evaporation. | TCV61 |
|  |  | **Bromeliads**  Presence  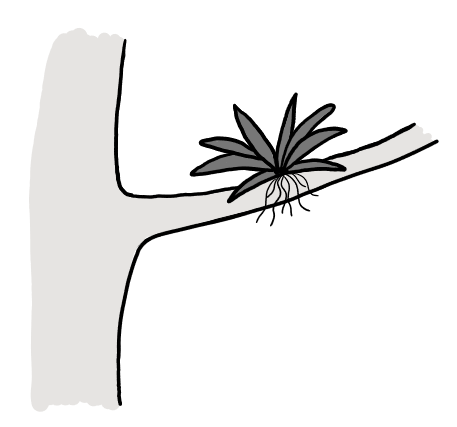 | Bromeliads growing directly on a part of the tree. | TCV62 |
| **The injuries and exposed wood** | Exposed sapwood only | **Bark loss**  Area > 300 cm²  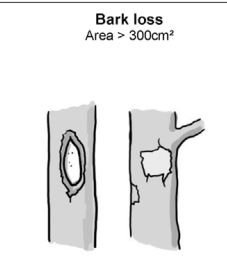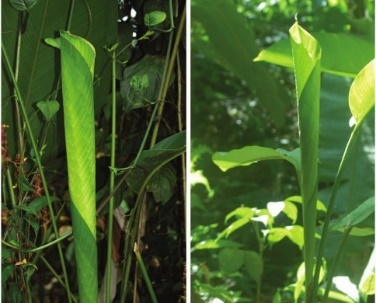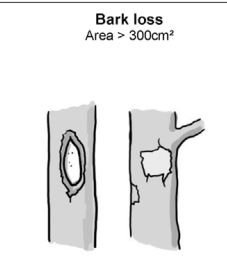 | Loss of bark exposing sapwood (skinning caused e.g. by felling, skidding, natural tree fall, rock fall, rodents). | TIN11 |
|  |  | **Fire scar**  Area > 600 cm²  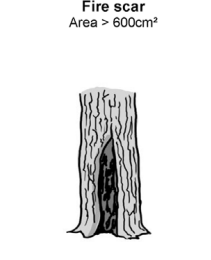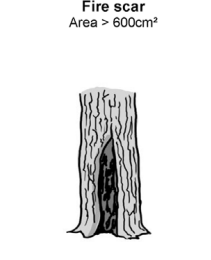 | Fire scars on the lower trunk. They usually have a triangular shape and are located at the base of the tree on the leeward side. Fire scars are associated with charcoal and sometimes resin flow on exposed sapwood or bark. | TIN12 |
|  |  | **Bark shelter**  Gap > 1 cm, depth > 10 cm, height > 10 cm  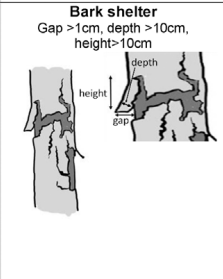  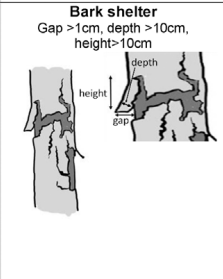 | Space between peeled-off bark and sapwood forming a shelter (open at the bottom). | TIN13 |
|  |  | **Bark pocket**  Gap > 1 cm, width > 10 cm, height > 10 cm  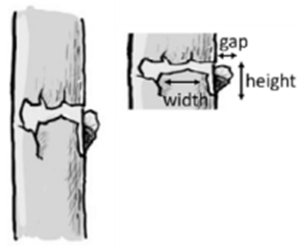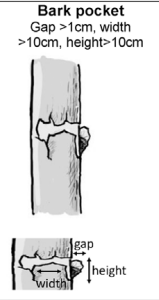 | Space between peeled-off bark and sapwood forming a pocket (open at the top) possibly containing mould. | TIN14 |
|  | Exposed sapwood and heartwood | **Stem breakage**  Ø > 10 cm at break point  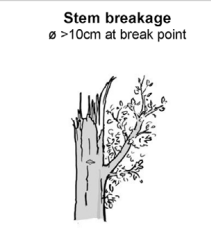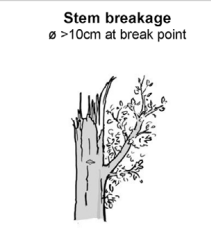 | The stem has broken off but the tree is still alive. The lower part of the deadwood is in contact with living wood with sap flow. | TIN21 |
|  |  | **Limb breakage**  Exposed heartwood  > 300 cm²  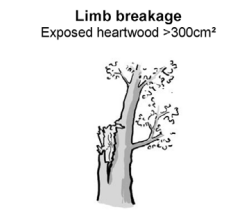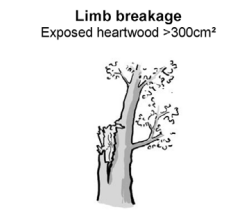 | Exposed heartwood through limb or fork breakage. The wound is surrounded by living wood with sap flow. | TIN22 |
|  |  | **Crack**  Length > 30 cm, width  > 1 cm, depth > 10 cm  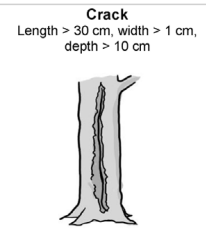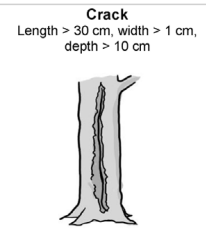 | Crack through the bark and the wood (if caused by lightning strike, see below). | TIN23 |
|  |  | **Lightning scar**  Length > 30 cm, width  > 1 cm, depth > 10 cm  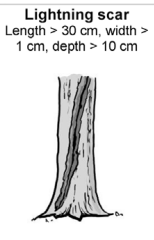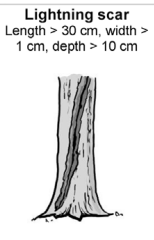 | Crack caused by lightning strike; usually spiraling around the tree with splintered wood present. | TIN24 |
|  |  | **Fork split at insertion**  Length > 30 cm  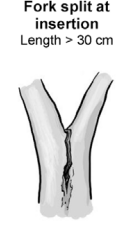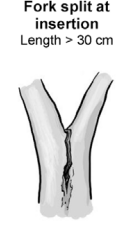 | Crack at the insertion of a fork. (If one side of the fork has broken off, see Stem breakage). | TIN25 |
| **Crown deadwood** | Crown deadwood | **Dead brances**  Branch Ø > 10 cm, or Branches Ø >3 cm and  > 10 % of the crown is dead  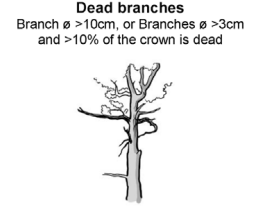 | Dead branches located in the canopy, conditions remain relatively shady. | TDE11 |
|  |  | **Dead top**  Ø > 10 cm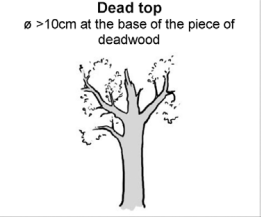 at the base of the piece or deadwood  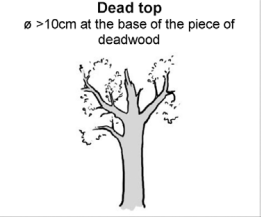 | The entire top of the tree is dead; the deadwood is sun-exposed | TDE12 |
|  |  | **Remaining broken limb**  Broken end Ø 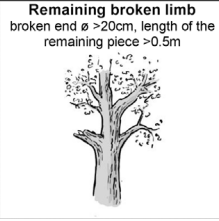> 20 cm, length of the remaining piece > 0.5 m  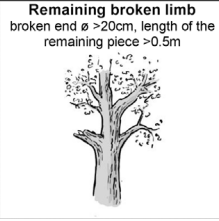 | A limb has broken off. The remaining end may be splintered. The injury does not affect the trunk (If so, see Stem breakage). | TDE13 |
| **Excerscences** | Twig tangles | **Witch broom**  Largest Ø > 50 cm  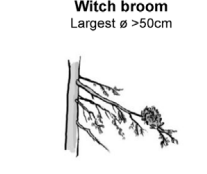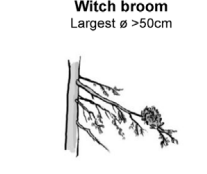 | Dense agglomeration of twigs on branches | TGR11 |
|  |  | **Epicormic shoots**  >5 twig clusters  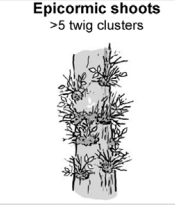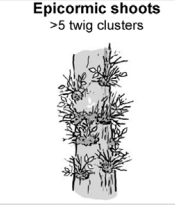 | Dense agglomeration of twigs along the trunk. | TGR12 |
|  | Burrs and cankers | 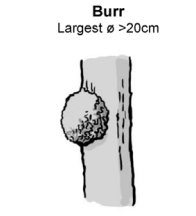**Burr**  Largest Ø > 20 cm  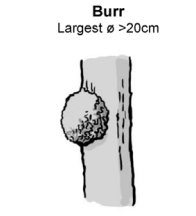 | Proliferation of cell growth with rough bark | TGR21 |
|  |  | 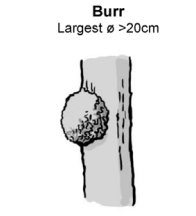**Canker**  Largest Ø > 20 cm or large part of the trunk covered  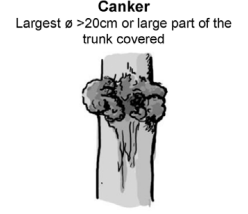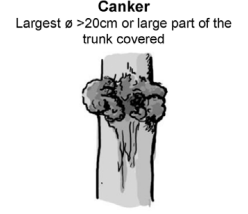 | Decayed canker. Sapwood exposed. Caused by e.g. Melampsorella caryophyllacerum, Nectria l. s. | TGR22 |
| **Fruiting bodies of saproxylic fungi and slime moulds** | Perennial fungal fruiting bodies | 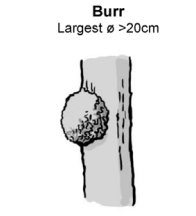**Perennial polypore**  Largest Ø > 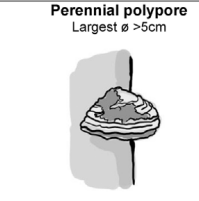5 cm  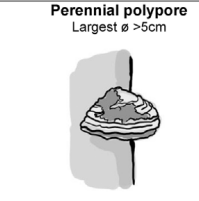 | Tough fruiting bodies of perennial polypores, showing distinct annual tube layers. | TFB11 |
|  | Ephemeral fungal fruiting bodies | 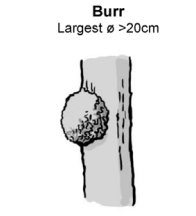**Annual polypore**  Largest Ø > 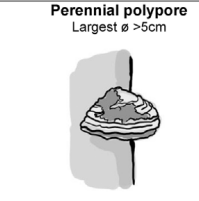5 cm or cluster of > 10 fruiting bodies  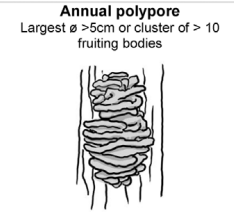 | Fruiting bodies of annual polypores, lasting several weeks. | TFB12 |
|  |  | 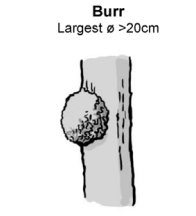**Pulpy agaric**  Largest Ø > 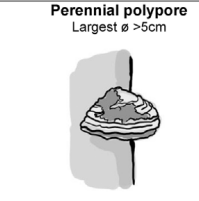5 cm or cluster of > 10 fruiting bodies  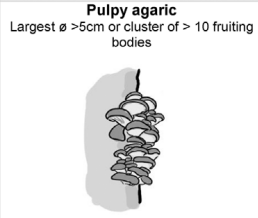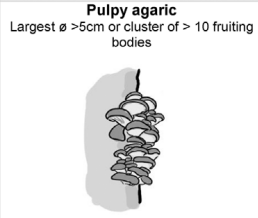 | Large, thick and pulpy or rather fleshy fruiting body of gillbearing fungi (order Agaricales). E.g.: Armillaria, Pleurotus, Pholiota, or large Pluteus species. The fruiting body generally remains several weeks. | TFB13 |
|  |  | 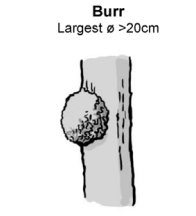**Large Pyrenomycete**  Stroma Ø > 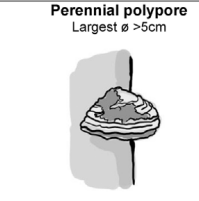3 cm or stroma cluster covering > 100 cm²  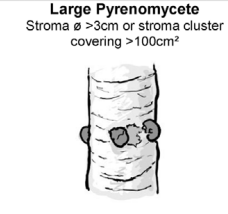 | Tough hemispheric dark fungi ressembling a lump of coal. E.g: Daldinia or Hypoxylon. | TFB14 |
|  |  | 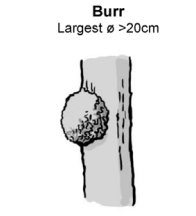**Myxomycetes**  Largest Ø > 5 cm  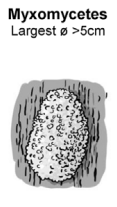 | Amoeboid slime mold which forms moving plasmodium. The plasmodium is gelatinous when fresh. | TFB15 |
| **Epiphyltic and epixylic structures** | Epiphytic and parasitic crypto- and phanerogams | 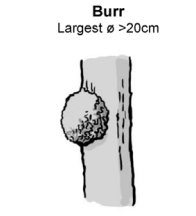**Bryophytes**  > 10 % of the trunk area covered  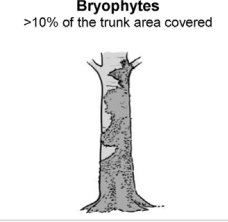 | Trunk covered by mosses and liverworts. | TEP11 |
|  |  | 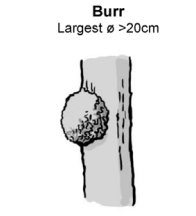**Foliose and fructicose lichens**  >10 % of the trunk area covered  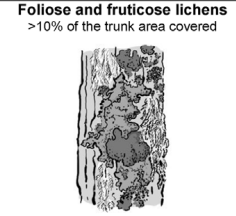 | Trunk covered by foliose or fruticose lichens. | TEP12 |
|  |  | **Ivy and lianas**  >10 % of the trunk area covered  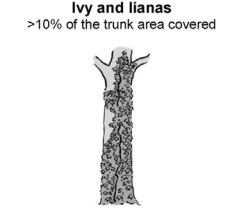 | Lianas and other climbing phanerogams (Hedera helix, Clematis vitalba, Lonicera periclimenum, Vitis vinifera). | TEP13 |
|  |  | **Ferns**  > 5 fronds  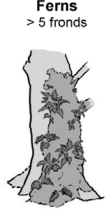 | Ferns growing directly on a part of a tree (i.e. epiphyte) | TEP14 |
|  |  | **Orchids**  Presence  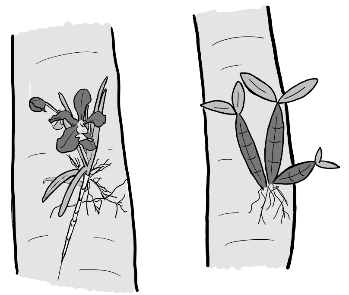 | Orchids growing directly on a part of a tree | TEP15 |
|  |  | **Mistletoe**  Largest Ø > 20 cm  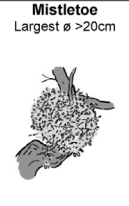 | Hemiparasitic plants (Viscum spp., Arceuthobium oxycedri, Loranthus europaeus). | TEP17 |
|  |  | **Hemiepyphyte**  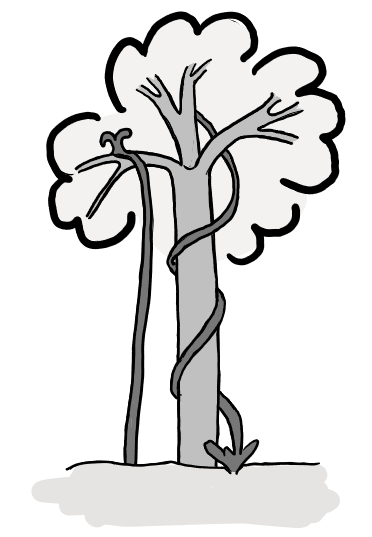 | Hemiepyphytes differ from epiphytes by having roots reaching the soil. | TEP18 |
|  |  | **Strangler Fig around living tree**  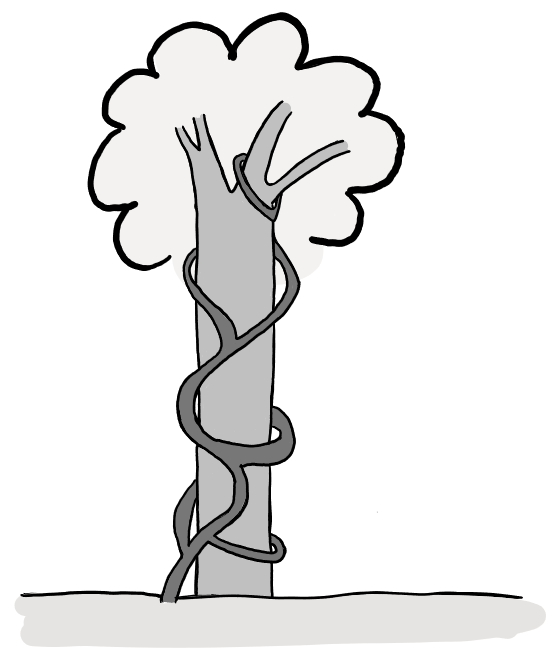 | Strangler Figs grow around trees and having roots reaching the soil. They can kill the inner tree and become the tree instead and profit from the rotting wood. | TEP19 |
|  |  | **Strangler Fig around dead tree**  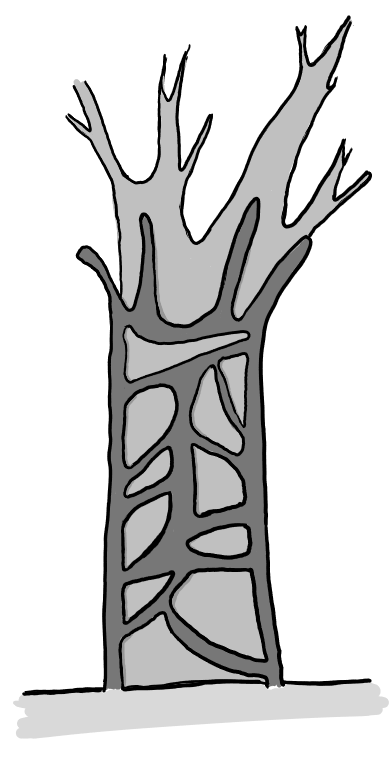 | Strangler Figs grow around trees and killed tree already | TEP110 |
|  |  | **Dead Lianas**  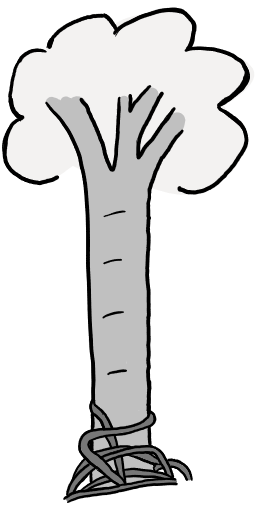 | Dead lianas fell of the trunk and aggregate around the trunk with ground contact. | TEP111 |
|  | Nests | **Vertebrate nest**  Ø > 10 cm  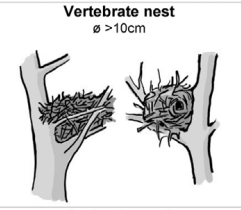 | Nest built by birds, dormice, mice or squirrels. | TEP21 |
|  |  | **Invertebrate nest**  Presence  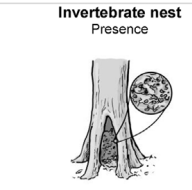 | Larval nest of invertebrates: e.g. Pine processionary moth Thaumetopoea pityocampa, wood ant Lasius fuliginosus or wild bees Apis mellifera. | TEP22 |
|  |  | **Cartonnest**  Presence  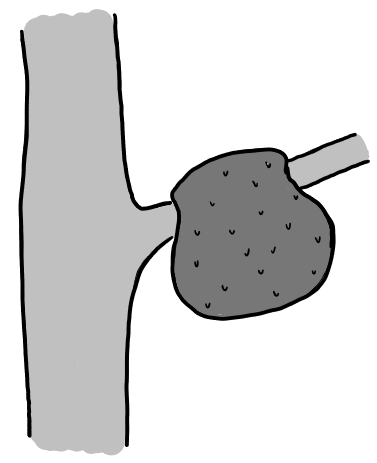 | Nests build by termites and ants on the tree. | TEP23 |
|  | Microsoils | **Bark microsoil**  Presence  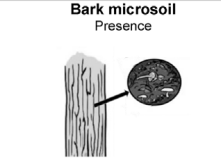 | Microsoil resulting from micro-pedogenesis of epiphytic mosses, lichens or algae and necrosed old, thick bark. | TEP31 |
|  |  | **Crown microsoil**  Presence  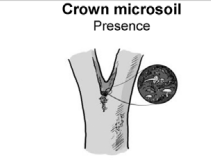 | Microsoil resulting from pedogenesis of debris and litter fallen from the crowns, often colonized by roots of the TreM bearingtree. Main positions: flat areas on limbs, forks, sometimes stem junctions of twin trees. | TEP32 |
|  |  | **Dead leaf**  >1 m  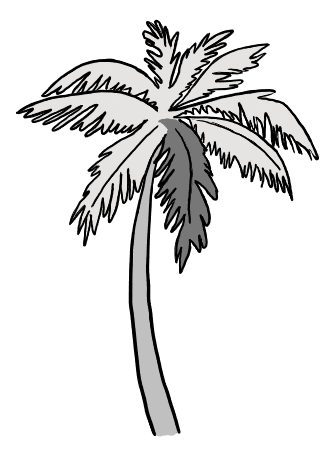 | Dead leaf still attached to the tree. Minimum size of 1 m. | TEP33 |
| **Exudates** | Exudates | **Sap run**  Cumulative length > 10 cm  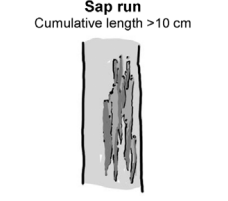 | Fresh significant flow of sap. | TOT11 |
|  |  | **Heavy resinosis**  Cumulative length > 10 cm  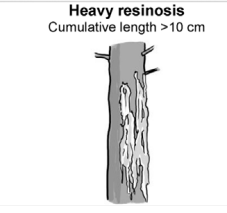 | Fresh significant flow of resin | TOT12 |
| **Root formations** | Root formations | **Stiltroots**  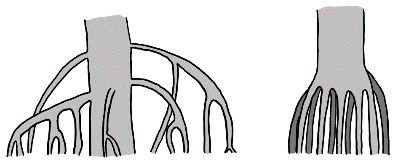  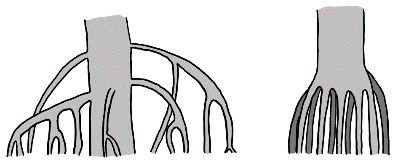 | Stilt roots arises from the lower bola and runs to the ground. In Mangroves and a few palms. | TRF11 |
|  |  | **Buttressroots**  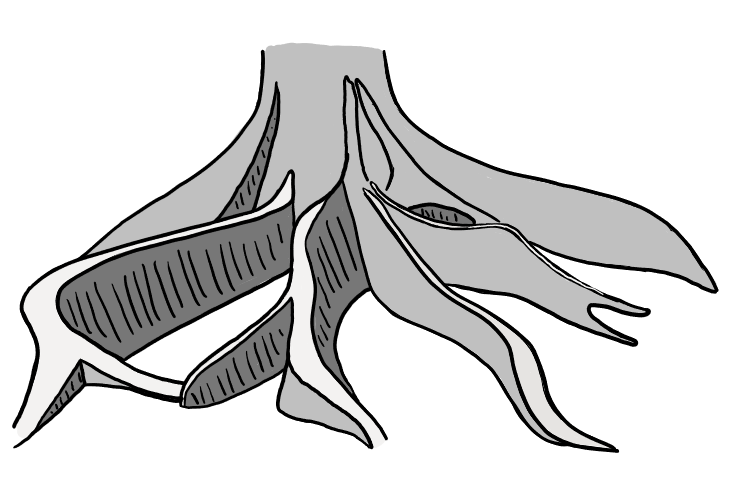 | Buttress roots are aearial extensions of lateral surface roots. | TRF12 |
